# Distinct perceptual and conceptual representations of natural actions along the lateral and dorsal visual streams: an EEG-fMRI fusion study

**DOI:** 10.1101/2025.09.17.676633

**Authors:** Diana C. Dima, Jody C. Culham, Yalda Mohsenzadeh

## Abstract

Actions are the building blocks of our dynamic visual world, yet the neural computations supporting action perception are not well understood. How does perceptual and conceptual information unfold in the brain when we observe what others are doing? We collected EEG and fMRI data while participants viewed short videos and sentences depicting naturalistic actions. Using representational similarity analysis, we found a posterior-to-anterior gradient along the lateral pathway, from perceptual features to conceptual, modality-invariant representations. Among conceptual features, the target of actions (i.e. whether the action was directed at an object, a person, or the self) explained the most unique variance in EEG responses. In fMRI, we found distinct conceptual representations along the ventral, dorsal, and lateral pathways, with the target of actions specifically encoded in lateral occipitotemporal cortex (LOTC) and posterior superior temporal sulcus (pSTS). Finally, EEG-fMRI fusion revealed rapid processing along the lateral and dorsal pathways. Together, our results disentangle the perceptual and conceptual components of action understanding and characterize the underlying spatiotemporal dynamics in the human brain.

## 1 Introduction

The visual world is always in motion. To make sense of what we see and decide on our own behaviours, we must understand what people are doing. Actions connect people, objects, and environments, and are thus a fundamental building block in our dynamic and adaptable perception. However, the sequence of neural computations that supports action understanding is not well understood. How does the brain extract perceptual and conceptual information from dynamic natural actions?

Actions vary along many perceptual axes (Leshinskaya et al., 2020; Lingnau and Downing, 2015), and can be defined in several ways. For example, an action may be described as a pattern of movement performed by particular body parts (effectors); as a group of behaviours that can be described using the same verb; or according to other essential characteristics, such as the use of a tool, the body part involved, the target of the action, etc. In previous behavioural work, we showed that people categorize naturalistic action videos and sentences according to the target of actions, that is, whether the action is performed upon an object, another person, or the self (Dima et al., 2024). Here, we used spatiotemporally resolved neural data to compare action representations in brain and behaviour and to characterize the processing of perceptual and conceptual action features over time along the lateral and dorsal visual streams.

By combining naturalistic stimuli, EEG and fMRI data, and theoretically informed models of action features, we addressed several gaps in the current understanding of action perception. First, we disentangled conceptual representations, including four different action taxonomies derived from prior work and a behaviourally driven model based on human similarity judgements (Dima et al., 2024), to understand *which* semantic information is represented in the brain during action perception. Second, we used EEG-fMRI fusion to understand the timing of processing in the action observation network, including along the lateral and dorsal visual streams. Finally, we used video and text renditions of everyday actions to understand how modality-invariant responses emerge during action perception.

Actions are thought to be processed rapidly, with semantic and invariant information emerging within 200 ms (Dima et al., 2022; Isik et al., 2018). The brain encodes actions at different levels of abstraction (Spunt et al., 2016; Wurm and Lingnau, 2015; Zhuang et al., 2023) and extracts invariant representations that generalize across visual characteristics (Hafri et al., 2017; Isik et al., 2018; Jastorff et al., 2010; Tucciarelli et al., 2015; Urgen and Orban, 2021) and even across vision and language (Aflalo et al., 2020; Wurm and Caramazza, 2019a). These representations span ventral, lateral, and dorsal areas in the action observation network (Caspers et al., 2010; Urgesi et al., 2014), whose roles in action perception remain the subject of debate. A lateral pathway spanning the lateral occipitotemporal cortex (LOTC) and posterior superior temporal sulcus (pSTS) is thought to process actions along a gradient from perceptual features (bodies, motion, tools, effectors; Beauchamp et al., 2003; Peelen et al., 2006; Wurm and Caramazza, 2019b) to conceptual information (invariant representations and social information; McMahon et al., 2023; Pitcher and Ungerleider, 2021; Tarhan and Konkle, 2020a; Wurm and Caramazza, 2022, 2019a), yet many studies report overlapping representations (Kabulska et al., 2024; Lingnau and Downing, 2015; Tucciarelli et al., 2019a). The ventral occipitotemporal cortex (VOTC) also encodes action features, particularly as they relate to scenes and objects (Chao et al., 1999; He et al., 2020; Wurm et al., 2017). The role of dorsal stream areas like the inferior parietal lobule (IPL) and premotor cortex remains unclear and has been hypothesized to relate to action goals or social affordances important in action planning (Culham and Valyear, 2006; Hamilton and Grafton, 2006; Orban et al., 2021). In addition, little is known about the timing of neural processing within the action observation network.

Here, we used a multimodal approach with naturalistic stimuli (Haxby et al., 2020; Nastase et al., 2020) and collected EEG and fMRI data while participants viewed short videos and sentences depicting everyday actions. We defined a set of 11 features ranging from perceptual (e.g. motion energy, number of agents, body parts involved) to conceptual (e.g. action class, action target, behavioural similarity) and investigated modality-invariant representations that generalize across vision and language. Our results reveal the rapid processing of actions and disentangle perceptual and conceptual representations along the lateral and dorsal visual streams.

## 2 Results

### 2.1 Decoding naturalistic actions from EEG and fMRI patterns

We collected EEG and fMRI data while participants (*N*=20 in each of two experiments; section 4.2) viewed 95 two-second videos depicting naturalistic actions from the Moments in Time dataset (Monfort et al., 2019). To understand the contributions of perceptual and conceptual information to action processing, the stimuli were annotated with features derived from image properties and experimenter-defined labels (section 4.4). Perceptual features ranged from low-level (motion energy and activations from the V1-like layer of CORnet-S) to features related to agents (number and gender) and actions (effectors and tool use). Conceptual features included the target of actions (object-, person- or self-directed), action class (based on a shared motor goal, e.g. locomotion or manipulation), everyday activity (based on common categories, e.g. sports or working), and action verb (the verb describing the action, e.g. running or welding).

These features defined actions at different levels of abstraction based on previous work (Dima et al., 2024; **Figure 1a**). A behavioural similarity feature was computed from a separate multiple arrangement experiment with the same stimuli (Dima et al., 2024; section 2.3). All features were converted to representational dissimilarity matrices (RDMs) encoding the distances between all pairs of videos. To investigate action representations across vision and language, participants also viewed 95 sentences describing the actions in each video in separate fMRI and EEG sessions (see section 4.1). The videos and sentences were presented in counterbalanced sessions while participants performed a one-back task on actions.

**Figure 1.**
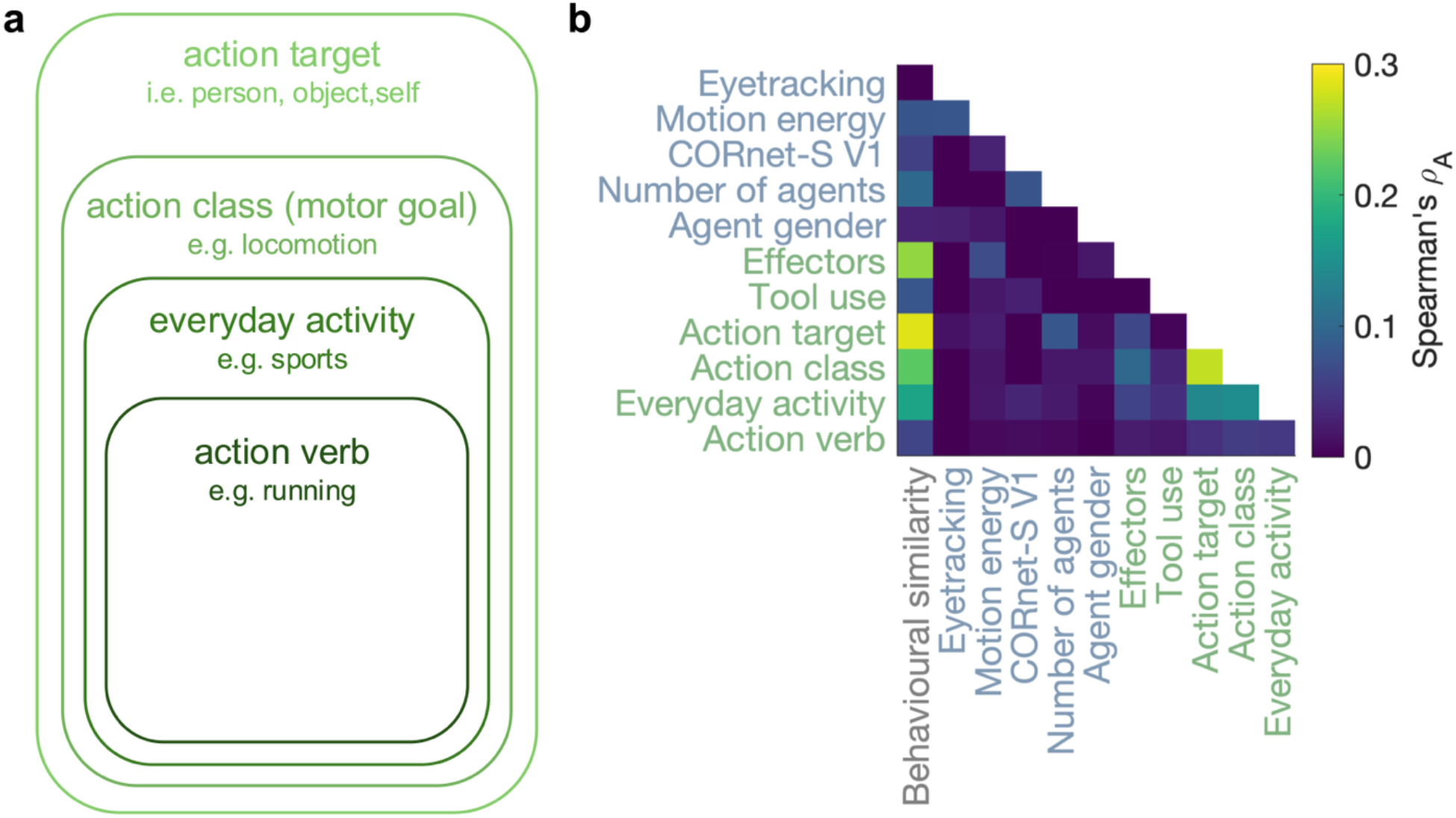
Stimulus features. **a**. Four semantic features captured different levels of abstraction within the action set. **b**. Correlations between feature RDMs in the video set, showing low correlations between perceptual and semantic features. As a control, a participant-averaged RDM of eyetracking data collected while participants viewed the videos (section 4.5) is included, and only correlates moderately with the motion energy RDM.

To characterize neural processing across space and time, we performed pairwise video and sentence decoding on fMRI and EEG responses. FMRI decoding was performed using 5-mm searchlights within areas with reliable responses determined using split-half reliability thresholding (Tarhan and Konkle, 2020b; section 4.6). EEG decoding was performed at each time point for a 2-ms temporal resolution.

In fMRI, videos were decodable in a broad range of visual areas along the ventral, lateral and dorsal streams (**Figure 2a**), with the highest accuracies obtained in left LOTC and left parahippocampal place area (PPA). EEG video decoding accuracies rose above chance at ∼57 ms after stimulus onset and remained so for the entire stimulus duration (**Figure 2b**). Sentences were decoded with lower accuracy in both datasets, mainly in early visual areas and between ∼120-780 ms after stimulus onset.

**Figure 2.**
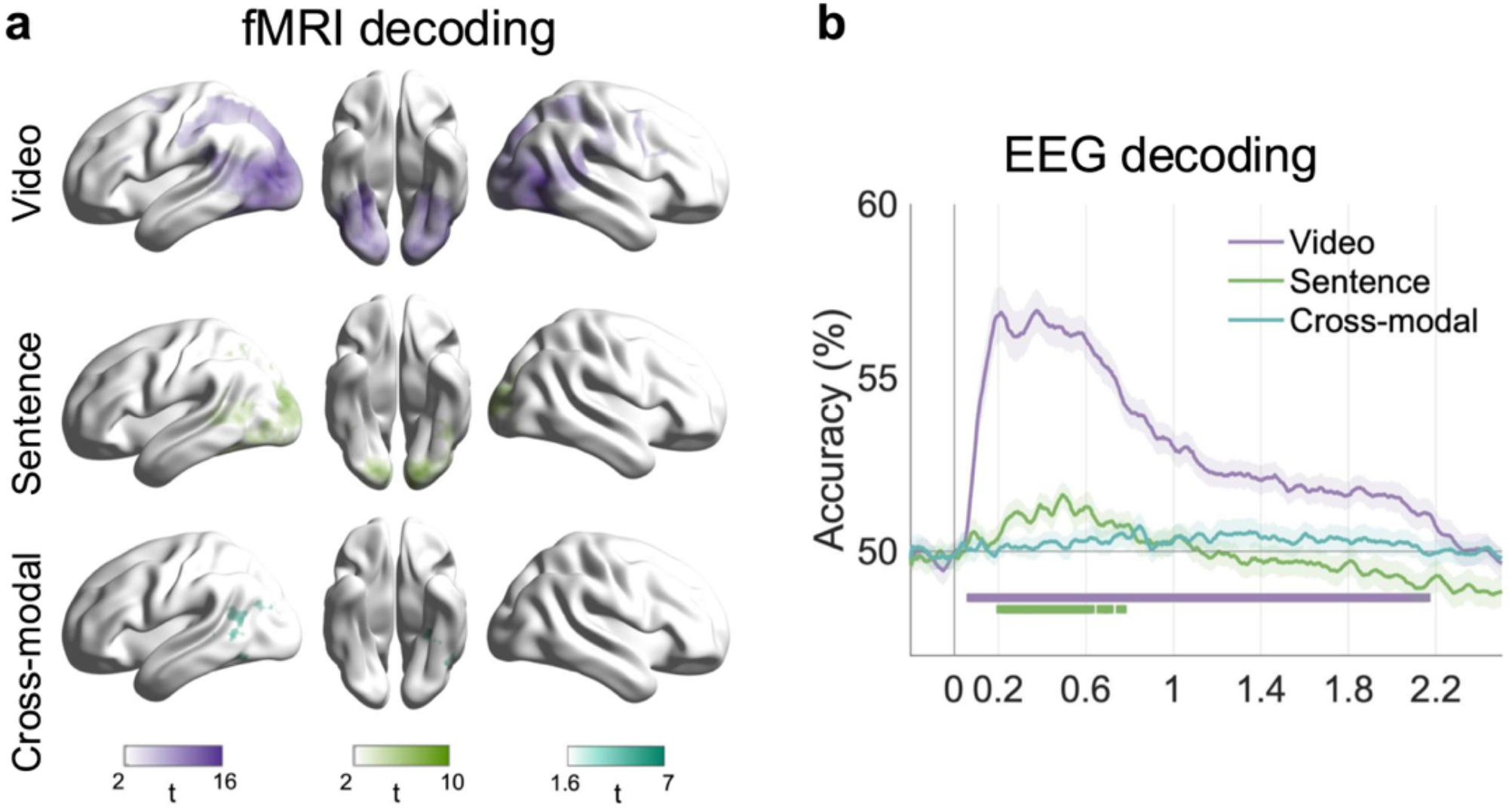
Decoding results for videos, sentences, and across modalities. Decoding was performed within modality (decoding all pairs of videos and sentences) and across modalities (by training a classifier on videos and testing it on sentences). **a**. Searchlight fMRI decoding results (t-values, TFCE-corrected *p*<0.05). **b**. Time-resolved participant-averaged EEG decoding accuracies. Significant time points are marked with horizontal lines and error bars are SEM.

To investigate modality-invariant representations, we trained and tested a classifier to decode each pair of actions across videos and sentences. In fMRI, cross-modality decoding reached above-chance accuracy mainly in a left LOTC cluster in the posterior middle temporal gyrus (MTG), extending into the inferior temporoparietal junction (TPJ). No statistically significant time points were detected in the EEG data.

### 2.2 How visual action information unfolds in the brain

Using representational similarity analysis, we investigated how action information unfolds in the brain, with a focus on disentangling different types of conceptual information. We used EEG decoding accuracies to compute time-resolved neural RDMs for each participant, and correlated them to the feature RDMs (Spearman’s *ρ*_*A*_). Motion energy correlated with the EEG patterns within 100 ms, while action effector information emerged after 180 ms, and semantic features after 200 ms (**Figure 3a**). Behavioural similarity correlated with the EEG patterns for most of the stimulus duration, starting at 120 ms (**Supplementary Figure 2**).

**Figure 3.**
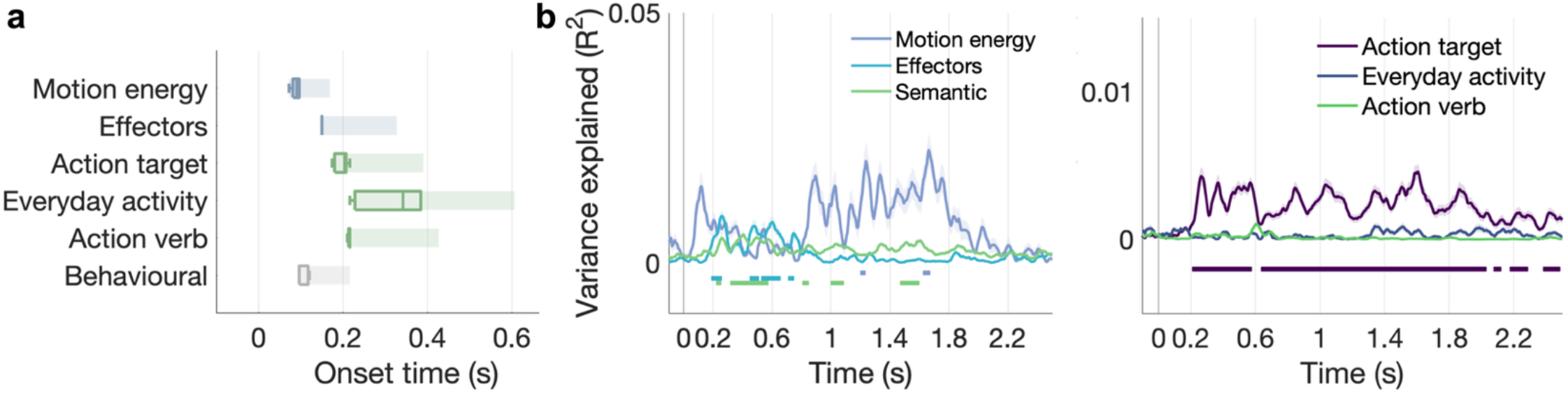
EEG feature representations for video stimuli. **a**. RSA analysis: RSA onset latencies for significant features. Boxplots show median onset latencies and rectangles show the associated 90% confidence intervals based on leave-one-subject-out onsets. **b**. Variance partitioning analysis: Unique variance explained by features in the variance partitioning analyses. In the left-hand panel, semantic features included the action target, action verb, action class, and everyday activity features. Solid lines show variance explained in the averaged neural patterns, and error bars show SEM across leave-one-subject-out iterations. Horizontal lines show significant time points across all leave-one-subject-out iterations.

To disentangle the unique contributions of perceptual and semantic features, we conducted two variance partitioning analyses (**Figure 3b**), using a hierarchical regression approach to assess the unique variance explained by different groups of features in the EEG data. First, we assessed the unique contributions of low-level visual features (motion energy), effectors, and the four semantic features. Results supported a perceptual-to-conceptual processing sequence, with semantic features explaining unique variance after 230 ms. Similar results emerged when including CORnet-S V1 activations as a visual feature (**Supplementary Figure 3**).

A further analysis disentangling the contributions of semantic features (action target, action verb, and everyday activity) revealed that only the action target explained unique variance in the EEG patterns from ∼200 ms, for most of the stimulus duration. A different grouping of semantic features yielded similar results (**Supplementary Figure 4**). Across both variance partitioning analyses, variance inflation factors were moderate, with a maximum of 1.83 (mean 1.35±0.29) and 1.32 (mean 1.22±0.09) respectively, suggesting a low risk of collinearity.

In the fMRI data, we used a searchlight multiple regression approach to assess the contribution of each feature RDM to the neural patterns while accounting for other features. Variance inflation factors ranged between 1.01 and 1.91, with a mean of 1.31±0.35. Among all features, the strongest representations were found for the behavioural similarity RDM across visual, lateral, and dorsal areas (**Figure 4**). Low-level features were represented in lateral and ventral visual cortex (CORnet-S), with motion energy representations extending anteriorly into the extrastriate body area (EBA) and pSTS. The number of agents was also represented in ventral and lateral areas extending into pSTS and TPJ, speaking to the perceptual and social information encoded by this feature.

**Figure 4.**
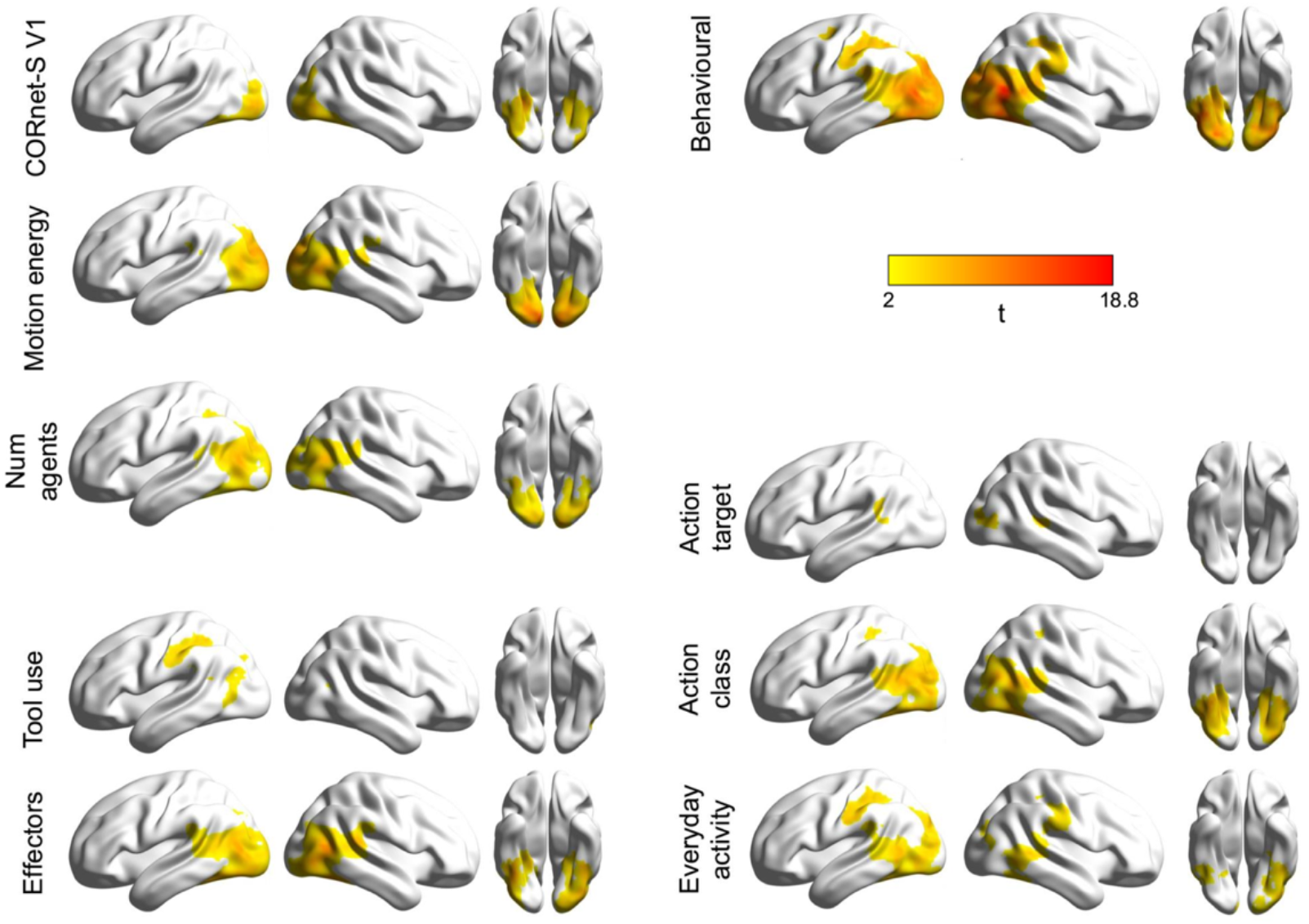
FMRI feature representations (multiple regression RSA) for video stimuli. Statistical maps show t-values corrected for multiple comparisons (TFCE *p*<0.05, 5000 permutations).

Representations of effectors spanned lateral and ventral areas, extending from LOTC into bilateral TPJ and ventrally along the fusiform gyrus. In line with previous work, tool use representations were mainly found in left IPS and anterior supramarginal gyrus, as well as left aITS.

Semantic features were represented in lateral, ventral, and dorsal areas. Along the lateral pathway, overlapping semantic representations were found in LOTC. Rather than effector-based representations, the dorsal stream housed action category representations. Specifically, action class was represented in LOTC, areas along the ITS and fusiform gyrus, as well as areas along the posterior occipital sulcus (POS) and intraparietal sulcus (IPS). The everyday activity feature spanned some of the same regions, particularly along bilateral pSTS and left IPS. The action target was represented in limited clusters in bilateral pSTS and right LOTC. There were no action verb representations after accounting for other features.

### 2.3 Spatiotemporal dynamics in action perception

Using EEG-fMRI fusion, we investigated action processing in space and time. We correlated EEG RDMs to participant-averaged fMRI RDMs extracted from 25 lateral and dorsal ROIs based on the Jülich cytoarchitectonic brain atlas (Amunts et al., 2020; **Figure 5a**). We found that V1 (hOc1) representations began to contribute to EEG patterns at ∼96 ms, while LOTC (hOc5, consistent with MT+; Malikovic et al., 2007) and TPJ emerged at ∼144 ms. Within the IPS, there was a temporal cascade from posterior regions such as hIP7 (retinotopic area IPS0; Richter et al., 2019) at ∼96 ms to anterior regions such as the anterior intraparietal area, aIPS; Choi et al., 2006) at ∼300 ms. Regions in the IPL emerged between ∼90-330 ms, again following a posterior-to-anterior cascade. Timing in the SPL diverged from the temporal gradients seen in IPS and IPL; specifically, activation was seen earliest in medial area 5 (5M and 5Ci), higher-order somatosensory cortex, after 190 ms before lateral area 5 (5L) and area 7 (7PC and 7M) after ∼300 ms. Searchlight EEG-fMRI fusion revealed widespread significant clusters in visual cortex within 80 ms of stimulus onset, spreading to higher-level regions within 200 ms (**Figure 5b; Supplementary Movie 1**).

**Figure 5.**
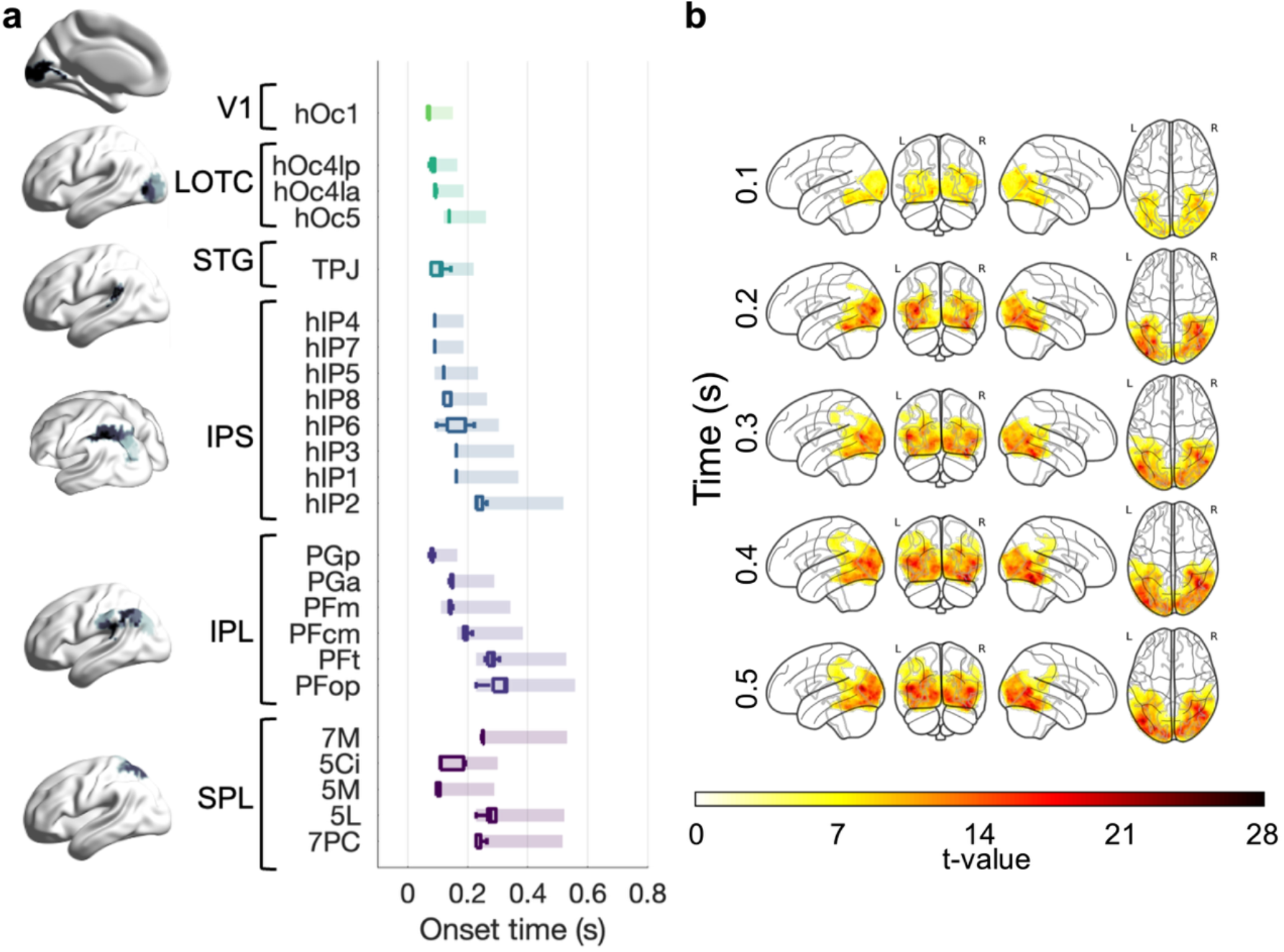
EEG-fMRI fusion for video stimuli. **a**. Onsets of correlations between EEG RDMs and fMRI RDMs extracted from left lateral and dorsal ROIs. Boxplots show medians and rectangles show 90% confidence intervals based on leave-one-subject-out onsets (jackknifing). Subregions are organized from posterior to anterior. LOTC: lateral occipito-temporal cortex, including hOc4lp (LO-2), hOc4la (LO-1), and hOc5 (MT+/V5); STG: superior temporal gyrus; IPS: intraparietal sulcus; IPL: inferior parietal lobule, from posterior (hIP7 or IPS0) to hIP3 (post-aIPS), hIP1 (vIPS), and hIP2 (aIPS); SPL: superior parietal lobule. **b**. Searchlight EEG-fMRI fusion results, shown in steps of 100 ms. Maps show significant t-values (TFCE *p*<0.05, 5000 permutations). See Supplementary Movie 1.

## 3 Discussion

We combined spatially and temporally resolved neural data with video and text stimuli to investigate how natural action perception unfolds in the brain, from perceptual information to conceptual representations. We report four main findings based on this work. First, action representations were consistently more robust for videos than for sentences, particularly in dorsal cortex, with cross-modal representations found mainly in anterior LOTC. Second, our results reveal a temporal gradient organizing action information from perceptual to conceptual, with the target of actions dominating EEG responses after 200 ms. Third, we find a perceptual-to-conceptual gradient along the lateral pathway and categorical representations of actions in dorsal areas. Finally, EEG-fMRI fusion reveals rapid processing (150 ms) in LOTC and relatively slower processing (within 350 ms) unfolding in a posterior-to-anterior cascade through parietal cortex, highlighting different temporal dynamics in the action observation network.

### 3.1 A temporal gradient of action information

We used four semantic features to define actions at different levels of abstraction, from action verbs to the target of actions, and disentangled their contributions to the neural patterns. In previous behavioural work, we found that the target of actions (object, person or self) best explained semantic similarity judgments of actions, independently of other semantic features like action class (Dima et al., 2024). Here, we report a similar pattern in the EEG data (**Figure 3**). In variance partitioning analyses, only the action target explained unique variance in the EEG data starting at ∼230 ms after stimulus onset (see also **Supplementary Figure 4**). This highlights the importance of action goals in action perception, linking action understanding and social inference (Blakemore and Decety, 2001; Spunt et al., 2011; Tarhan et al., 2021; Thornton et al., 2019).

The latency of this effect is in line with previous work suggesting that semantic action information emerges within ∼200 ms (Dima et al., 2022; Isik et al., 2018). Although effect sizes were low, semantic features explained variance in the EEG data independently of perceptual information, including action effectors. Our results reveal a perceptual-to-conceptual gradient, from low-level visual features extracted within 100 ms to perceptual action information (effectors) and semantic information within 200 ms.

### 3.2 Conceptual information along the lateral pathway

FMRI results also revealed a gradient of representations in LOTC (**Figure 4**), from low-level visual features and motion energy in posterior areas, to more conceptual information in anterior areas. Similarly to Wurm and Caramazza (2019a), we found modality-invariant action representations in left anterior LOTC. This is a striking result given differences in stimulus sets (eight types of hand actions depicted via controlled stimuli, versus 95 perceptually varied videos and sentences sampling a large action set). The lateralization of the effect is in line with work showing that semantic representations are restricted to left LOTC after controlling for perceptual features (Tucciarelli et al., 2019a; Wurm and Caramazza, 2019b). This does not preclude the existence of conceptual representations in other areas, such as the modality-invariant response to action verbs previously reported in parietal cortex using single cell recordings (Aflalo et al., 2020); however, these representations may serve different goals (Wurm and Caramazza, 2022).

The semantic representations of action class, everyday activity, and action target overlapped, but there were also important distinctions between them. Action class and everyday activity representations also extended to ventral areas and dorsally along the IPS, in line with suggestions that behaviourally relevant action categories are represented along the dorsal stream, enabling computations relevant in action planning (Orban et al., 2021; Orban et al., 2021; Urgen and Orban, 2021). On the other hand, the action target was mainly represented in lateral areas, particularly in TPJ and right LOTC. Growing evidence suggests that social information is processed in pSTS, with some proposals suggesting a social processing goal for the lateral pathway (Isik et al., 2017; McMahon et al., 2023b; McMahon and Isik, 2023; Pitcher and

Ungerleider, 2021). The target of actions reflects social information (i.e. whether an action is directed towards an object, the self, or other people). Thus, these results align with previous evidence that TPJ performs agent detection during action observation (Wurm and Schubotz, 2018), and more generally with the growing evidence of the importance of social features in action perception (Dima et al., 2022; Tarhan and Konkle, 2020a; Wurm et al., 2017). On the other hand, we find a more restricted and anterior representation of the action target compared to previous work after accounting for other conceptual information like the action class.

The anterior location of these representations is in line with a posterior-to-anterior gradient enabling the extraction of conceptual and social information along the lateral pathway.

Furthermore, EEG-fMRI fusion results revealed LOTC and TPJ processing within 150 ms after stimulus onset, suggesting that this information is computed in a rapid feedforward sweep. The lateral pathway has been proposed to process visual motion (Boussaoud et al., 1990), multimodal information (Weiner and Grill-Spector, 2013), action representations (Lingnau and Downing, 2015; Wurm and Caramazza, 2022), and dynamic social perception (Pitcher and Ungerleider, 2021). Our results are in line with the latter proposals, suggesting that conceptual and socially relevant action information is processed along the lateral stream.

We also found widespread and overlapping representations of intermediate features. In line with previous work, the number of agents was strongly represented in LOTC and VOTC (Papeo, 2020; Wurm and Caramazza, 2019b), but also extended dorsally along the IPS. Effectors were represented in ventral and lateral areas, including into pSTS, potentially pointing to their role as precursors for semantic and social information. Tools were represented in left LOTC and IPS, in line with previous work highlighting parietal encoding of tool use planning and observation (Culham and Valyear, 2006; Leshinskaya et al., 2020; Peeters et al., 2013). No representations of action verbs emerged in the multiple regression RSA analysis, potentially due to the low reliability of our dataset in anterior regions along the lateral pathway, where action verb representations have been reported (Bedny et al., 2008; Peelen et al., 2012).

### 3.3 Dorsal representations of conceptual information

While past research has often emphasized the roles of the lateral and ventral streams in action perception, our results also provide novel insights about dorsal stream representations. Within the dorsal stream, neural representations (particularly in the left IPS, postcentral sulcus, anterior SMG, and premotor cortex) were explained by behavioural similarity, but not by perceptual features. Consistent with earlier evidence that behavioural similarity reflects conceptual action information (Dima et al., 2024), the dorsal stream also represented semantic features such as action classes and everyday activity categories. Interestingly, semantic features were more aligned with dorsal stream representations than action effectors in the multiple regression RSA analysis. A long-standing debate surrounds dorsal stream representations, with the classic textbook view arguing for a somatotopic representation of effectors (Penfield and Boldrey, 1937), and the ethological view arguing for the primacy of action goals and a common organizations across primate species (Graziano and Aflalo, 2007; Stepniewska and Kaas, 2023). While studies have found effector-specific activation in parietal cortex (Buccino et al., 2001), there is growing evidence of action-based representations in the dorsal stream (Jastorff et al., 2012; Lorey et al., 2014). Our data provide new evidence that action goals may be more relevant than effectors for the dorsal stream when using a naturalistic range of everyday actions and multivariate analyses that include both effectors and action features. This information about action goals may support mentalizing computations in STS (Spunt and Lieberman, 2012; Thornton and Mitchell, 2018). While the processing of tool-use actions in the lateral and dorsal streams, especially in anterior SMG, is well established (Ishibashi et al., 2016; Lewis, 2006; Orban and Caruana, 2014; Reynaud et al., 2016), our results suggest that the dorsal stream also represents the broad spectrum of everyday actions.

Dorsal stream processing for simple sensory-motor tasks is typically thought to be fast due to magnocellular geniculostriate pathway inputs (Schmolesky et al., 1998). However, EEG-fMRI fusion reveals relatively slow onset latencies in higher-level dorsal regions in the IPL and SPL compared to lateral regions like motion-selective MT+/V5 and LOTC regions associated with body and object perception. Although anterior parietal regions are thought to play an important role in action observation (Molenberghs et al., 2012), our results show that their representations are among the latest to emerge, following a posterior-to-anterior cascade from more posterior parietal regions and even from higher-order somatosensory regions (area 5), which perhaps have faster access to information via multisensory links between sensory cortices. Taken together, these results raise interesting questions about the potential reliance of dorsal stream regions on ventral and lateral stream representations, as well as somatosensory representations during action observation.

Naturalistic videos and sentences straddle the boundary between actions and events, and our findings may thus provide a window into the processing of real-world events and natural vision more broadly. Together, our results characterize the neural dynamics of natural action perception in space and time, from perceptual features to conceptual representations that generalize across vision and language. These findings open exciting avenues for understanding the ventral, lateral, and dorsal stream computations supporting dynamic vision.

## 4 Methods

### 4.1 Stimuli

The stimulus set consisted of 95 short videos depicting everyday actions (**Figure 1**). Videos were selected from the Moments in Time dataset (Monfort et al., 2019), resized to 400 × 600 pixels, trimmed to 2 s, and resampled to 20 frames/s (see Dima et al., 2024). In addition to the experimental set, 20 different “catch” videos were included as targets for a one-back task (10 pairs of stimuli depicting the same action).

To assess modality-invariant action representations, each video was accompanied by a sentence describing the action, in the format Agent + Action [+Object, if present] + Context, e.g. “Two men are talking in a living room.” Sentences were presented in black Arial font on a white 400 × 600 pixel background with gray borders to match the video stimuli.

### 4.2 Participants

Behavioural data were collected in a previous online multiple arrangement experiment (Dima et al., 2024), with the final sample including 39 participants (age 18.1±0.8, 22 female, 16 male, 1 non-binary). Participants were recruited through the Department of Psychology Research Participation Pool at Western University.

A sample of 20 participants took part in the EEG experiment (age 24±3.2, 13 female, 6 male, 1 non-binary), and 20 participants took part in the fMRI experiment (24.5±5.3, 14 female, 6 male). Six participants took part in both experiments. All participants were right-handed, had normal or corrected-to-normal vision, and reported no history of neurological or psychiatric disorders.

Recruitment was conducted through the OurBrainsCAN participant pool at Western University using online announcements and flyers, and participants were compensated for their participation. All procedures for online data collection were approved by the Western University Research Ethics Board, and informed consent was obtained from all participants.

### 4.3 Behavioural experiment

In the behavioural experiment (Dima et al., 2024), participants arranged the videos according to the semantic similarity (i.e., the similarity in meaning) of the depicted actions. The experiment was conducted via the Meadows platform (https://meadows-research.com/) and entailed arranging different subsets of videos to reach sufficiently robust distance estimates for each pair (Kriegeskorte and Mur, 2012). Inverse MDS was applied to generate representational dissimilarity matrices from participants’ arrangements as a proxy for the semantic organization of actions in the mind (Dima et al., 2024).

### 4.4 Feature definitions

To understand how perceptual and conceptual information is processed in the brain while viewing natural action videos, we quantified a set of visual and semantic features based on image properties and experimenter annotations of the stimulus set. Visual features were converted to representational dissimilarity matrices (RDMs) by computing the pairwise Euclidean distances between feature vectors corresponding to each video. Semantic features were operationalized as binary RDMs where distances were set to 0 between similar videos and to 1 between dissimilar videos.

To capture visual features, motion energy was computed for each video using a pyramid of spatiotemporal Gabor filters with the *pymoten* package in Python 3.10.9. Additionally, activations were extracted from the V1-like layer (V1 Conv2) of the biologically-inspired CORnet-S model pretrained on ImageNet (Kubilius et al., 2019, 2018). Agent information was captured using two experimenter-labeled features: the number of agents (one, two, three or more) and the perceived gender of agents. Two additional features were created to quantify perceptual features related to actions, i.e. effectors and the use of tools. The effector feature characterized each video with a six-element binary vector indicating the involvement of different body parts in the action (face/head, hands, arms, legs, torso). The binary tool use feature indicated whether the action depicted in each video was tool-mediated or not.

Four semantic features categorized the stimuli at different levels of abstraction based on previous work (Dima et al., 2024; see **Supplementary Table 1**). The action target feature grouped actions into three categories: object-directed, person-directed, or self-directed. The action class feature grouped actions into eight categories based on a shared motor goal, such as locomotion, ingestion and manipulation (Orban et al., 2021). The everyday activity feature included activity categories inspired by the American Time Use Survey (ATUS, 2019) frequently used in action research. Finally, the action verb feature grouped videos characterized by the same verb label in the Moments in Time dataset.

In addition, a behavioural similarity feature was derived from the multiple arrangement data (section 2.3) by averaging the behavioural RDMs across participants.

### 4.5 EEG experiment

#### 4.5.1. Experimental procedure

Participants viewed the videos and sentences in two separate EEG sessions on different days, whose order was counterbalanced across participants. In each session, continuous EEG data were recorded using a 64-channel BioSemi system with a sampling rate of 1024 Hz. Stimuli were presented on a monitor with a 60 Hz refresh rate, and a photodiode was used to accurately track stimulus presentation times. Eyetracking data were collected during each EEG recording using an EyeLink Duo eyetracker with a sampling resolution of 1000 Hz.

The 95 stimuli were presented in pseudorandom order in 10 blocks separated by self-paced breaks. To maintain attention, participants performed a one-back task on actions and pressed a button when the same action was depicted in two consecutive videos or sentences. Catch stimuli (10 pairs) were randomly presented and repeated twice throughout the experiment. To minimize learning effects, repeated catch pairs were presented in different orders. In total, each session consisted of 990 trials (950 experimental trials and 40 catch trials).

Stimuli were presented for 2 s and separated by a variable inter-stimulus interval (ISI) chosen from a uniform distribution between 1 and 1.5 s. A black fixation cross was displayed on a grey background during the ISI. Stimuli were displayed on the same grey background, and the fixation cross remained on screen during video presentation, but not during the sentences. Participants were instructed to fixate when viewing the videos. The paradigm was implemented in MATLAB R2021a using the Psychophysics Toolbox (Brainard, 1997; Kleiner et al., 2007; Pelli, 1997).

#### 4.5.2 Preprocessing

EEG data were preprocessed using MATLAB R2020b and the FieldTrip toolbox (Oostenveld et al., 2011). First, the EEG data were aligned to stimulus onset using the photodiode data to correct for any lag between stimulus triggers and on-screen presentation. Next, the data were segmented into 3 s epochs (0.2 s pre-stimulus to 2.8 s post-stimulus onset), baseline-corrected using the 0.2 s prior to stimulus onset, and high-pass filtered at 0.1 Hz.

A semi-automated pipeline was applied for artifact rejection. To detect muscle artifacts, the data were filtered between 110 and 140 Hz and Hilber-transformed before removing segments with a z-value over 15. High-variance channels and trials were manually rejected after visual inspection using the *ft_rejectvisual* function in FieldTrip to generate summary plots. Eye movement components were identified and removed with independent component analysis (ICA). Catch trials and trials that included a button press were removed from the data (7.04±2.8% of trials on average). After artifact rejection, a further 12.6±5.9% of trials were removed, and an average of 2.9±2.5 noisy EEG channels were removed from 15 datasets. Prior to analysis, the data were resampled to 500 Hz and low-pass filtered at 30 Hz to investigate evoked responses.

Eyetracking time-series data were segmented into epochs (2 s after stimulus onset), missing segments and blinks were removed, and time-resolved gaze positions (vertical and horizontal) were extracted. As a control in the video data analysis, eyetracking RDMs were created by calculating the pairwise Euclidean distances between the time-resolved gaze position vectors corresponding to each video.

#### 4.5.3 Time-resolved decoding

To characterize the temporal dynamics of neural processing, time-resolved decoding was performed using a linear support machine classifier (LibSVM, Chang and Lin, 2011). At each time point, the classifier was trained to decode every pair of stimuli from the whole-brain EEG data using two-fold cross-validation. The procedure was repeated ten times, and classifier accuracy was averaged across iterations and folds. Prior to classification, the data were standardized by the standard deviation of the training set and whitened using the covariance matrix of the training data (Guggenmos et al., 2018).

The analysis was performed separately for videos and sentences. Additionally, to investigate modality-invariant neural representations, a cross-decoding analysis was performed in which the classifier was trained on each pair of videos and tested on the corresponding pair of sentences at each time point.

#### 4.5.4. Time-resolved representational similarity analysis

Video decoding accuracies were used as dissimilarity measures to generate time-resolved neural RDMs. Using representational similarity analysis, each participant’s neural RDM was correlated with the feature RDMs to investigate visual and semantic representations in action perception.

The analysis was performed using 10 ms sliding windows with a 4 ms overlap, and Spearman’s *ρ*_*A*_ was used as a correlation metric to ensure differences in model complexity do not bias results (Schütt et al., 2023). A noise ceiling was calculated by correlating each participant’s neural RDM to the average RDM of all other participants (lower bound; Nili et al., 2014).

To further account for correlations between features, we performed a variance partitioning analysis. Feature RDMs were entered as predictors in a hierarchical regression to quantify the unique variance (R^2^) explained by each feature in the participant-averaged neural RDM. Using a leave-one-out approach, the analysis was repeated with subsamples excluding each participant to calculate effect onsets (see below). This approach was used to disentangle the contributions of (1) different semantic features and (2) perceptual and semantic features.

#### 4.5.5 Statistical testing

Statistical significance was assessed using a jackknife-based baseline criterion approach (Kiesel et al., 2008; Lahner et al., 2024; Miller et al., 1998) to address the low signal-to-noise ratio and issues with estimating latencies using cluster approaches (Sassenhagen and Draschkow, 2019). For multivariate analyses, the standard deviation of baseline values (the 200 ms immediately before stimulus onset) was computed based on the average timecourse across subjects. Next, a jackknife approach was applied to compute leave-one-subject-out timecourses. For each subsample, a time point was considered significant: (1) if its value was greater than or equal to twice the baseline standard deviation; and (2) if the average value across the next 10 windows of 50 samples (decoding) or 5 windows of 5 samples (RSA) was also greater than or equal to twice the baseline standard deviation. Onset latencies were calculated based on the first significant time point across all leave-one-out subsamples.

### 4.6 fMRI experiment

#### 4.6.1 Experimental procedure

Participants viewed the videos and sentences in separate fMRI sessions, counterbalanced across participants. Stimuli were presented in a rapid event-related design in 10 runs, each consisting of 119 trials and lasting approximately six minutes. Each run included 95 experimental trials, 20 null trials, and 4 catch trials presented in pseudsrandom order. In the experimental and catch trials, videos were shown for 2 seconds and followed by a one-second fixation cross on a gray background. In the null trials, a fixation cross was shown for 3 seconds. A fixation cross was presented for 6 seconds at the end of each run. Participants performed a one-back task on actions as in the EEG experiment.

FMRI data were acquired on a Siemens 3-Tesla Magnetom Prisma Fit MRI scanner using a T2*-weighted, single-shot, GE-EPI sequence (TR: 1000 ms; TE: 30 ms; flip angle: 48°;

FOV: 210 mm; multiband factor: 5; slice thickness: 2.5 mm; voxel size: 2.5 mm isotropic; 60 axial slices covering the whole brain). Structural images were acquired halfway through each session using a T1-weighted MPRAGE sequence (TR: 2300 ms; TE: 2.98 ms; flip angle: 9°; FOV: 256 mm; slice thickness: 1 mm; voxel size: 1 mm isotropic; 192 sagittal slices).

#### 4.6.2 Preprocessing

FMRI data were preprocessed using fMRIPrep version 20.2.7. Anatomical images underwent brain extraction, tissue segmentation, and spatial normalization to the MNI152NLin2009cAsym template. Functional images were slice-timing corrected, susceptibility-derived-distortion-corrected, and coregistered to the anatomical images. The preprocessing steps were applied using ANTs (http://stnava.github.io/ANTs/), FSL166, and FreeSurfer (http://surfer.nmr.mgh.harvard.edu/).

Next, single-trial responses were extracted using GLMsingle (Prince et al., 2022). Briefly, this approach fits an optimal haemodynamic response function at each voxel, denoises the data using GLMdenoise (Kay et al., 2013), and applies custom regularization of the betas at each voxel using fractional ridge regression. All analyses were conducted in MNI space, separately for each participant and session. No spatial smoothing was applied.

#### 4.6.3 Reliability thresholding

We evaluated the reliability of the video session data and performed voxel selection by computing the split-half reliability across odd and even runs for each participant and stimulus (Tarhan and Konkle, 2020b). Grand-average reliability maps were then liberally thresholded at the 90^th^ percentile (*r*>0.068; **Supplementary Figure 1**). Further analyses were performed within this reliability mask. To ensure consistency, the same mask was applied to the sentence session data, which had lower reliability.

#### 4.6.4 Searchlight decoding

Similarly to the EEG analysis, a linear classifier was used to decode each pair of videos and sentences, as well as across modalities (by training it on videos and testing it on sentences). Decoding was performed using a searchlight approach within the reliability mask. Searchlights had a radius of 5 mm (93 voxels total).

#### 4.6.5 Searchlight representational similarity analysis

To perform representational similarity analysis, neural RDMs were created at each searchlight location. Betas were z-scored across videos within each run and across voxels within each searchlight before computing the squared Euclidean distance between all pairs of videos. To account for correlations between models, we applied a multiple regression RSA approach, in which all models were entered as predictors (Tucciarelli et al., 2019b; Zhuang et al., 2023).

#### 4.6.6 Statistical testing

To assess the significance of accuracy and correlation maps, one-sample, one-tailed t-tests against chance and 0 respectively were conducted at each searchlight. T-values across searchlights were compared with a maximal statistic distribution obtained through permutation testing (5000 iterations), and the resulting p-values were corrected using a threshold-free cluster enhancement (TFCE) approach (α=0.05; Nichols and Holmes, 2001; Smith and Nichols, 2009).

### 4.7 EEG-fMRI fusion

EEG-fMRI fusion was conducted to resolve neural patterns in space and time (Cichy et al., 2016; Cichy and Oliva, 2020; Mohsenzadeh et al., 2019, 2018). In brief, this method correlates temporally resolved EEG patterns with spatially resolved fMRI patterns to characterize the flow of neural information processing.

First, a region-of-interest (ROI) approach was applied to determine the temporal dynamics of lateral and dorsal representations. RDMs from 25 left lateral and dorsal ROIs (**Figure 5**) capturing areas with reliable fMRI responses were extracted using maximum probability maps from the Jülich atlas (Amunts et al., 2020). Searchlight RDMs (section 2.6.5) were averaged within each ROI and across participants. The fMRI ROI RDMs were then correlated with the time-resolved neural RDMs using 10 ms sliding windows with 4 ms overlap (section 2.5.4).

Significance and onset latencies were determined using a jackknifing baseline criterion approach (section 2.5.5).

Next, to more broadly characterize the spatiotemporal dynamics of action processing, we correlated each fMRI searchlight RDM within the reliability mask with EEG RDMs averaged across 20 ms time windows. The resulting correlations were statistically assessed at each time window using TFCE-corrected one-sample t-tests (α=0.001, 5000 permutations).

