## Supplementary for "Distinct perceptual and conceptual representations of natural actions along the lateral and dorsal visual streams: an EEG-fMRI fusion study"

| Action verb | Everyday activity | Action class | Action target |
| --- | --- | --- | --- |
| applauding | childcare | defense | object |
| arresting | cooking | ingestion | person |
| assembling | eating | locomotion | self |
| baking | fighting | manipulation | |
| barbecuing | gardening | self-directed | |
| bicycling | grooming | social: gesture | |
| biting | hiking | social: interaction | |
| boating | instructing | social: symbolic | |
| boxing | painting | | |
| brushing | shopping | | |
| climbing | socializing | | |
| combing | sports | | |
| crafting | transportation | | |
| drinking | walking | | |
| driving | working | | |
| drying | writing | | |
| eating | | | |
| fencing | | | |
| fighting | | | |
| gardening | | | |
| hiking | | | |
| hitting | | | |
| hugging | | | |
| kicking | | | |
| painting | | | |
| picking | | | |
| reaching | | | |
| running | | | |
| stretching | | | |
| swimming | | | |
| talking | | | |
| walking | | | |
| washing | | | |
| waving | | | |
| working | | | |
| writing | | | |

**Supplementary Table 1**. Semantic categories in the stimulus set.

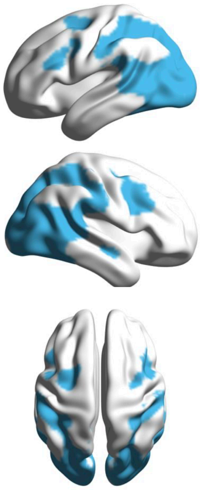

**Supplementary Figure 1**. Voxel reliability mask used in fMRI analyses.

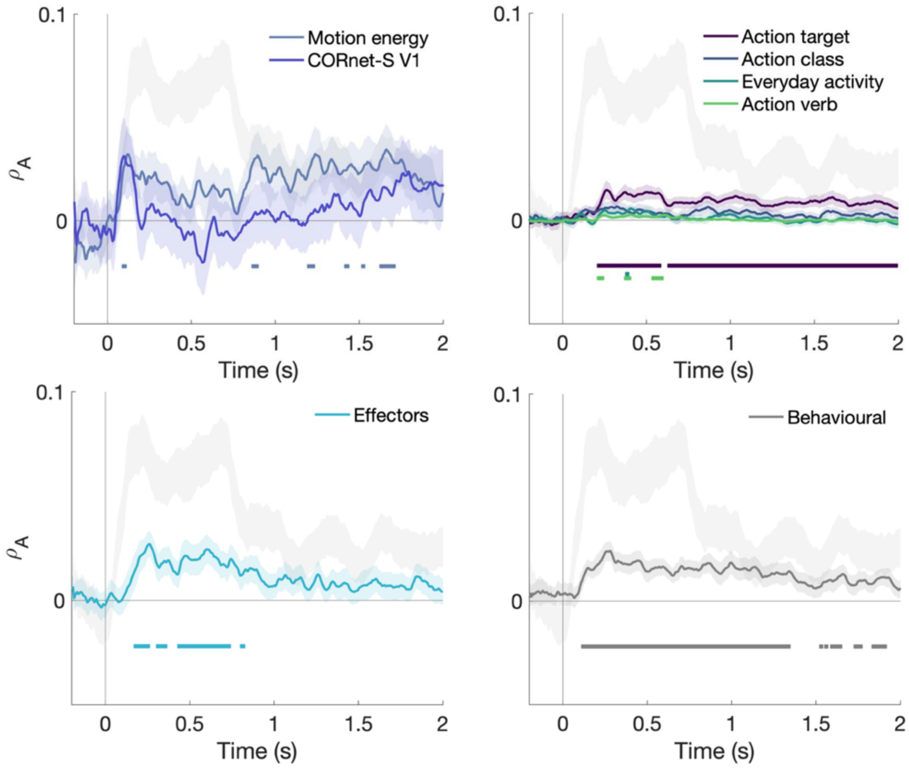


**Supplementary Figure 2**. Time-resolved RSA. Plots show the time-resolved correlation ($\rho_A$) of each significant feature RDM with the EEG data (error bars are SEM across participants). Significant time windows are marked with horizontal lines (jackknifing baseline criterion). The noise ceiling (mean+SEM) is shown in gray.

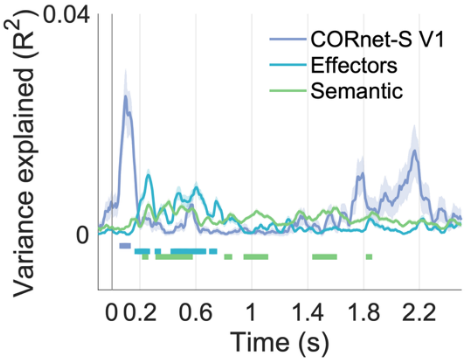


**Supplementary Figure 3**. Unique variance in EEG patterns explained by visual features (CORnet-S V1), effectors, and semantic features (action target, class, verb, and everyday activity) in a variance partitioning analysis. Solid lines show variance explained in the averaged neural patterns, and error bars show SEM across leave-one-subject-out iterations. Horizontal lines show significant time points across all leave-one-subject-out iterations.

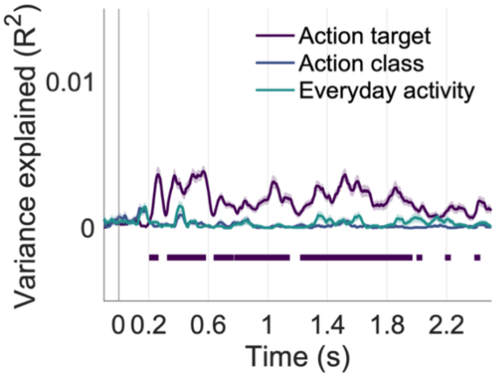


**Supplementary Figure 4**. Unique variance in EEG patterns explained by semantic features in a variance partitioning analysis. Solid lines show variance explained in the averaged neural patterns, and error bars show SEM across leave-one-subject-out iterations. Horizontal lines show significant time points across all leave-one-subject-out iterations.
